# Namibian fairy circles: Hostile territory for soil nematodes

**DOI:** 10.1101/2024.12.04.626864

**Authors:** Amy M. Treonis, L. Andrew Bell, Eugene Marais, Gillian Maggs-Kölling

## Abstract

Fairy circles are rings of grass with centers of bare soil that are found in some arid grasslands. Above- and belowground chemical and biological attributes of fairy circles have been explored in an ongoing debate about the ultimate causes of this pattern. We studied the soil nematode communities associated with *Stipagrostis* fairy circles along a 900-km range in the Namib Desert of Namibia in southern Africa. Nematode abundance and diversity were highest in soils along the vegetation rings that define fairy circles and in soils in the vegetated matrix surrounding the bare circles, demonstrating the positive impact of plant-derived resources on nematode communities. In contrast, soils from the bare centers of fairy circles had lower organic matter content and were nearly defaunated, averaging only 9.9 ± 1.7 nematodes 100 g^−1^ soil. Bacterial-feeding *Acrobeloides* nematodes were the only taxa over-represented in center soils in comparison to ring soils. Our results indicate that nematode communities are influenced by the unique soil environments that the fairy circle vegetation pattern generates and suggest that the soils at the centers of fairy circles are uniquely hostile habitat for soil organisms as well as plants. Co-occurrence network analysis of nematode communities elucidated relationships among the taxa. For example, the abundances of dorylaims and *Nothacrobeles* were positively correlated across all soil positions, suggesting they have overlapping ecological niches. In ring soils, the abundances of fungal-feeding *Aphelenchoides*, *Ditylenchus*, and *Hexatylus* were positively correlated, likely due to enhanced fungal communities in these plant-influenced soils. *Panagrobelus* demonstrated niche specialization by being negatively correlated to two other bacterial-feeding taxa (*Elaphonema* in matrix soils and *Acrobeloides* in ring soils). The co-occurrence patterns revealed by these relatively low diversity communities provide insights into the potential roles of nematode interactions as well as environmental factors in community assembly.

## Introduction

Fairy circles are a unique and enigmatic vegetation pattern found in some arid grasslands, originally reported from Namibia [1–4], but also found in Australia [5] and elsewhere in the world [6]. This pattern consists of over-dispersed, bare circular patches, 2-12 m diameter, that are each surrounded by a ring of grass that may have enhanced growth relative to the vegetation between the rings (Fig 1). In southern Africa, fairy circles form in windblown, sandy soils with 50-150 mm annual precipitation and occur along a range extending from Angola, through Namibia, to South Africa [2–4, 7]. This area is the transition zone between the sparsely vegetated gravel plains or sand dunes of the Namib Desert to the west and the arid shrub savanna of the interior. The grasses associated with fairy circle occurrences generally are perennial species of *Stipagrostis* [4, 7] that provide forage for wildlife, including zebra, oryx, springbok, and ostrich.

**Fig. 1.**
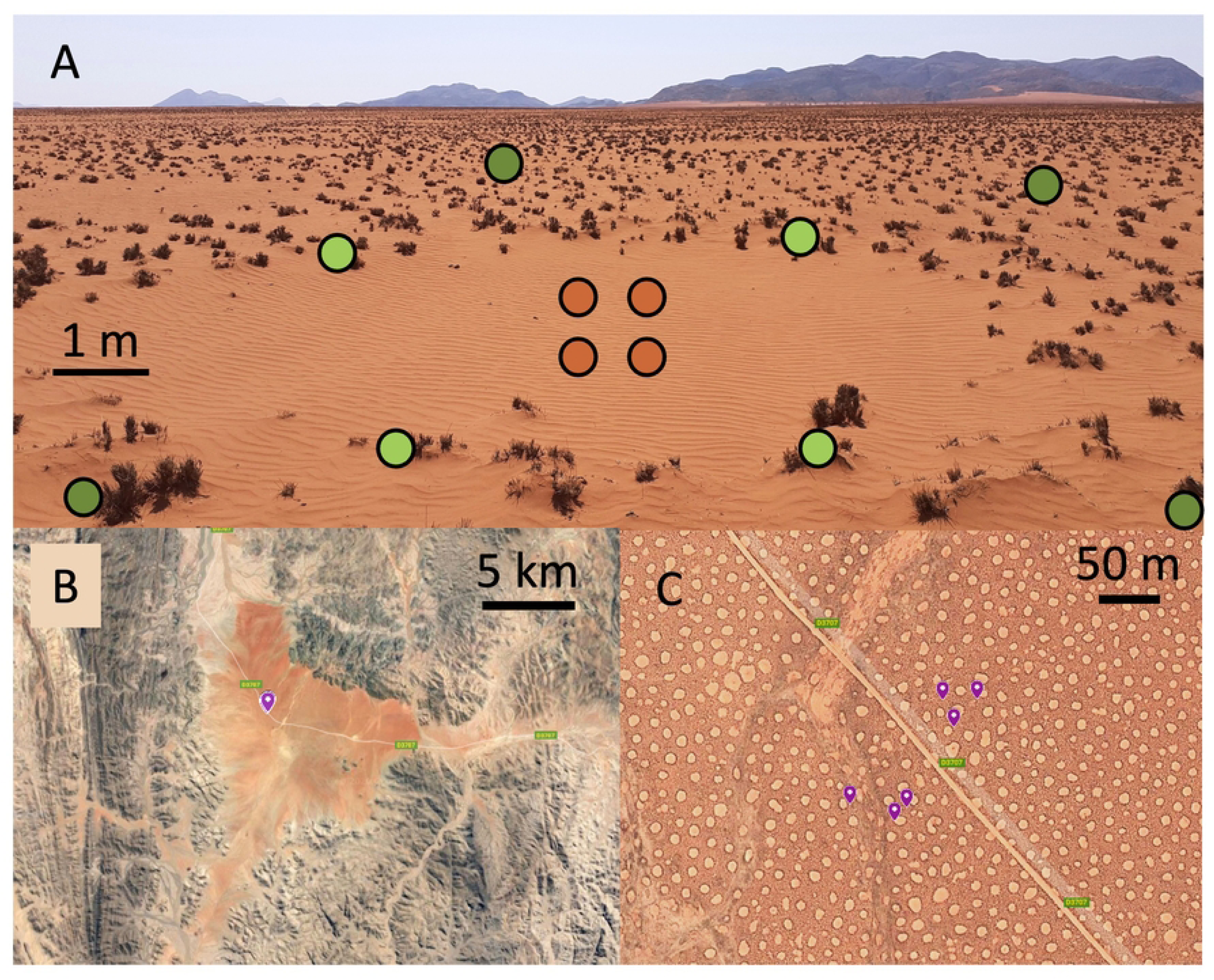
Fairy circles in the Giribes Plains, Kunene region, northwest Namibia. (A) A fairy circle. Brown dots represent sampling in the “center”, light green dots in the “ring”, and dark green dots in the surrounding “matrix”. (B) & (C) Satellite views with fairy circle sampling sites marked in purple [8].

The mechanism through which fairy circles form continues to be a topic of intense debate and study [4, 6, 9–11]. Various processes have been considered, including radioactivity [7], toxic gas leaks [12, 13], *Euphorbia* shrub allelopathy [14, 15], and microbial pathogens [16–18], with vegetation self-organization and insect herbivory as the two leading explanations. One hypothesis is that *Stipagrostis* grasses self-organize into geometric ring patterns in response to belowground competition for water and nutrients [4, 11, 19–22]. Alternatively, it has been proposed that the activity of ants or subterranean termites creates the bare patches due to foraging and/or to ensure that rainwater percolates into deeper soil layers [2, 9, 10, 23–27]. A logical conclusion seems that intersecting ecological phenomena play roles in fairy circle formation [6] and maintenance [9, 28].

So far, studies of the biological and hydrological properties of fairy circles have failed to consider soil nematodes. As members of the soil food web, these microscopic roundworms contribute to decomposition and nutrient cycling in soils [29] as they rely on plant-derived resources either directly (as plant-parasites) or indirectly (as consumers of decomposer bacteria and fungi) [30]. Soil nematode communities are very sensitive to vegetation patterns and plant diversity [31–34]. Hence, nematodes may provide insights into how soil habitability is affected in fairy circle landscapes. The presence of fairy circles influences soil organic matter, particle size, pH, salinity, moisture content, and temperature [2, 18, 22, 28, 35–37]. Bacterial and fungal communities have been shown to respond to this patterning, with distinct communities found in center, ring, and matrix soils [16–18]. Soil nematodes may be similarly responsive.

Abiotic and plant-related factors have often been seen to be the main drivers of nematode species distributions [38]. Frameworks to study soil nematode communities conventionally focus on the impacts of disturbance or organic enrichment, and taxa have been designated as r- or K-selected based on how they respond to environmental changes [39, 40]. These frameworks have been powerful for understanding the relationships between abiotic environmental factors, organic matter, and nematode communities [41–43]. However, they reveal little about the influence of nematode:nematode interactions and niche specialization in forming communities. Consequently, we lack an integrated understanding of nematode biodiversity patterns [44]. Recent studies demonstrate that belowground biotic interactions can contribute to nematode community assembly [45, 46]. Low-diversity soil nematode communities in arid environments provide a unique opportunity to study species interactions due to their relative simplicity [45, 47].

The objective of this study was to investigate the relationship between the fairy circle vegetation pattern and soil nematode diversity and abundance. We hypothesized that nematode abundance would vary among positions within the fairy circles (centers, rings, and surrounding matrix). Nematode community structure is influenced by the distribution of plant-derived resources, and communities in the ring and matrix soils should be more similar than either is to communities in the center soils. We also employed a co-occurrence network analysis approach to explore the role of species interactions in nematode community assembly and to attempt to identify assemblages of co-occurring nematode taxa.

## Materials and methods

### Study Sites

Nine fairy circle sites were sampled, spanning a 900-km north-south range that encompasses most of the distribution of fairy circles in Namibia [4, 7] (Fig 2). Eight sites were along the eastern verge of the Namib Desert, and the remaining site (Namib Naukluft) was within the most arid core of the desert. Due to drought, there was little fresh plant growth at any of the sites, and most of the grass was grazed to short stubs (Fig 1). At some sites (Namib Rand, Tsiseb), a small number of forbs and other grasses were present in the matrix area between fairy circles. During non-drought periods, plants grow in the rings and throughout the matrix, while the centers of the fairy circles remain devoid of vegetation [4] (Fig 1). It was not possible to identify the *Stipagrostis* species present due to the lack of growth. Soils were sandy Eutric Regosols (Tsiseb, Farm Bloemhof, Rostock, Namib Rand) or Petric Calcisols (Marienfluss, Giribes, Twyfelfontein, Namib Naukluft, Tsondab). Soils were collected under a research permit issued to A. Treonis (RPIV01022019) by the Namibia National Commission on Research Science and Technology. Permission to work in sampled areas was provided by the Namibian Ministry of Environment, Forestry and Tourism (Namib Naukluft, Tsondab Valley), the Namib Rand Nature Reserve (Namib Rand), private landowners (Farm Bloemhof, Rostock), and the communal conservancies.

**Fig 2.**
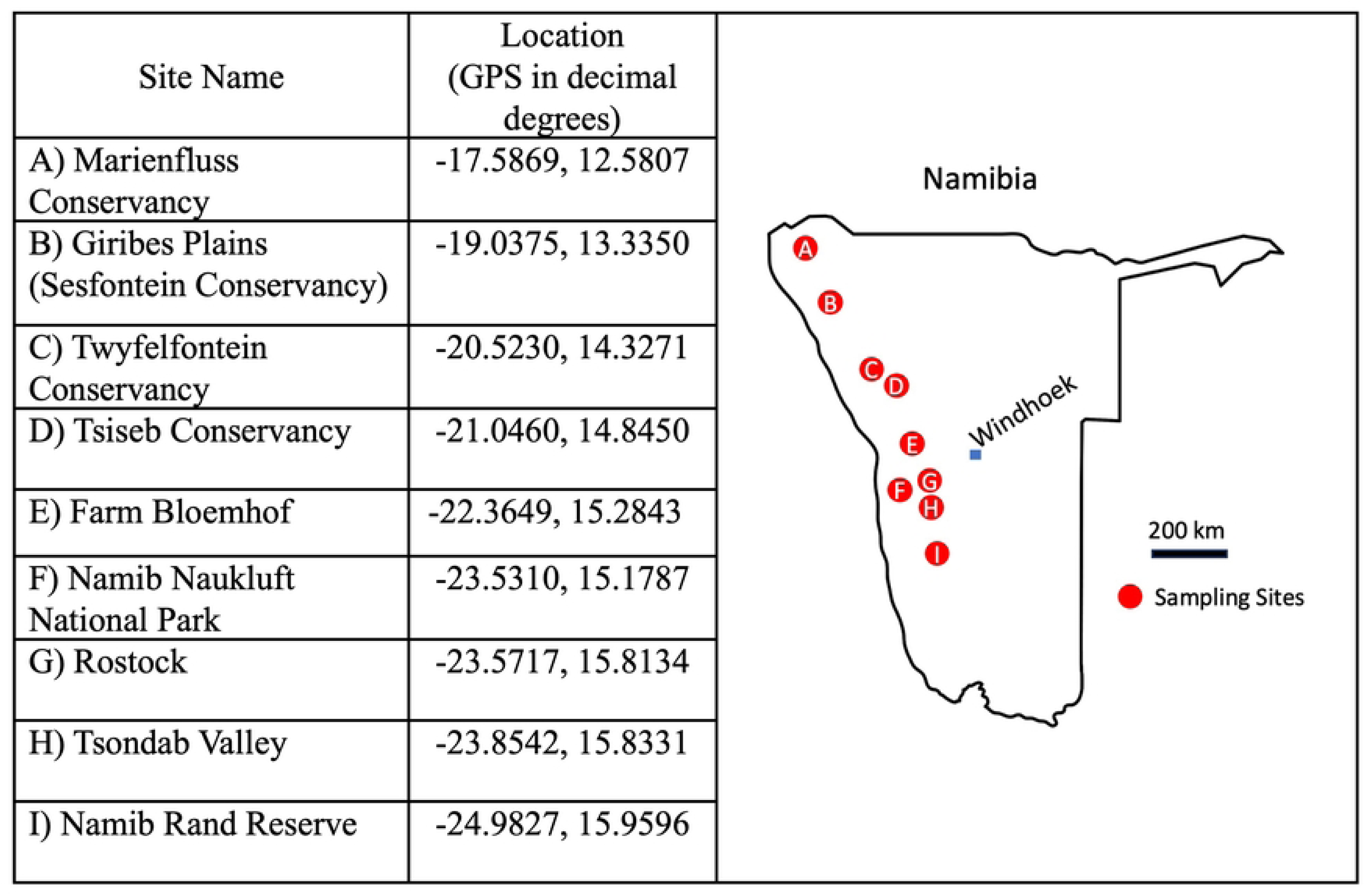
Namib Desert fairy circle sites that were sampled for study of soil nematode communities.

### Soil Sampling

Soils were collected between 24 March – 31 October 2020. Due to travel restrictions during the Covid-19 pandemic, sampling could not be completed within a short time frame. However, due to severe drought conditions, very little rainfall occurred in the Namib Desert over the sampling period [48], thus nematodes in these soils would have mostly been inactive throughout the study period (i.e., anhydrobiotic and not feeding or reproducing), and it is not likely that community composition would have changed.

Six representative circles were selected at each site within an approximately 1 ha area. At each circle, four subsamples were collected and combined from the center of the circle (“center”) to a depth of 0-10 cm using a plastic scoop (Fig 1). Similarly, four subsamples were collected and combined from the soil beneath the grass composing the ring of vegetation defining the margin of the fairy circle (“ring”), and four subsamples were collected and combined from the inter-circle matrix (“matrix”), at least 1 m from the ring (Fig 1). Matrix samples were collected from beneath vegetation, but the plants tended to be smaller than those defining in the ring. A total of 162 samples were collected (nine sites x six rings x three positions). Soils were sieved (2-mm mesh) to remove rocks and transferred to plastic bags. The dry, sandy soil contained very few rocks and passed easily through the sieve, minimizing any physical impact on nematodes.

### Soil Analyses

Soil moisture was determined gravimetrically (24 h at 105°C [49]). Soil organic matter was measured as loss on ignition (LOI, 360°C, 2 h [50]). Suspensions of 10-g air-dried soil in 30-ml deionized H_2_O were mixed and equilibrated for 30 min before solution pH was measured with a Pocket Pro+ pH Tester (Hach, Loveland, CO). Using the same solution, electrical conductivity was measured as an indicator of salinity using a conductivity meter (Traceable^®^, Cole-Parmer, Vernon Hills, IL [48]).

### Nematode Community Analyses

Nematodes were extracted over 72 h from 120 g soil using a Baermann funnel technique (60 g soil x 2 funnels per sample, combined when drawing off the solution[51]). Sample volume was reduced to approximately 500 µl, and nematodes were fixed in 5% hot:cold formalin solution. Nematodes were counted and identified to the lowest taxonomic level possible using a Zeiss inverted microscope (Carl Zeiss, Inc., White Plains, NY). Most nematodes were identified to the genus level, although certain groups with very low abundance and that were only found as juveniles could not be confidently identified at low taxonomic levels (i.e., dorylaims and plant-parasites in the family Dolichodoridae). References consulted for nematode identification include Bongers [52] and Andrássy [53]. Nematode taxa were assigned to trophic groups according to Yeates et al. [30]. Nematode abundance data were used to calculate nematode density (# 100 g^−1^ soil) and the Simpson’s Diversity Index [54]. The proportional representation of bacterial-feeding nematodes in the microbivore community was calculated as B/(B+F), where B was the number of bacterial-feeders and F was the number of fungal-feeders in the sample.

### Statistical Analyses

Statistical analyses were performed with R version 4.4 (https://www.r-project.org) [55]. The Shapiro-Wilk test was used to assess all dependent variables for normality. Analysis of variance (ANOVA) was used to investigate differences in soil organic matter content and electrical conductivity (log-transformed) among positions in the fairy circles across the sites. Means were compared using Tukey’s Honestly Significant Difference (HSD) multiple comparison procedure. Variables that could not be normalized by transformation (nematode abundance, soil moisture, pH, B/(B+F), Simpson’s Index) were analyzed with a Kruskal-Wallis nonparametric test followed by Dunn’s test for multiple comparisons, with *P*-values adjusted with a Bonferroni correction. Spearman’s correlation coefficient was calculated to investigate relationships among variables.

Nematode community structure among sites and fairy circle positions was explored via permutational multivariate analysis of variance (PERMANOVA) applied to Bray-Curtis distance, based on 999 permutations, and using the adonis2 function of the *vegan* package [56]. A Bray-Curtis based nonmetric multidimensional scaling (NMDS) approach was used to visualize the degree of similarity among nematode communities among fairy circle positions and sites. A constrained linear canonical ordination (RDA, [57]) was performed using the R *vegan* package [56] to test the contribution of field site, fairy circle position, and standardized soil properties in explaining the abundances of nematode taxa, which were Hellinger-transformed prior to RDA analysis. The RDA model was built using forward selection of variables, which result in the exclusion of soil moisture and organic matter from the analysis in favor of variables that explained a significant proportion of the variation. Five samples that did not contain any nematodes (one sample each from Rostock, Tsondab, and Farm Bloemhof, and two from Tsiseb; all center samples) and six taxa that were represented by ten or fewer nematodes across the study were excluded from multivariate analyses.

We used a co-occurrence network analysis approach to investigate connections among taxa within communities, excluding taxa represented by ten or fewer nematodes. Thresholds of 0.3 (positive correlations) and −0.2 (negative correlations) (all *P* < 0.05) were applied to filter for meaningful correlations from the Spearman’s correlation matrix of taxa abundances, resulting in an adjacency matrix that served as the basis for our network graph. An undirected graph was built using the Python NetworkX package, where nodes represent nematode taxa and edges indicate significant correlations. Centrality measures were computed to quantify the importance of each taxon within the network. Degree centrality was used to determine the node size, reflecting the number of direct connections each taxon had. Betweenness centrality was calculated for edges to highlight key connections that act as bridges in the network, with edge thickness proportional to betweenness centrality values. Closeness centrality was used to color nodes, indicating how centrally located a taxon is within the network. Community detection was performed using the Girvan-Newman algorithm [58] to identify clusters of species that interact more frequently with each other than with those outside their community.

## Results

### Soil Properties

Across the study sites, soils were very dry with low organic matter content (Table 1). Soil from the ring and matrix contained more organic matter than soil from the centers of the fairy circles (ANOVA, p=0.001, Table 1). Electrical conductivity (EC) was lowest in the center soils, followed by the matrix and highest in the ring (ANOVA, p<0.001, Table 1). Moisture and pH did not differ among fairy circle positions (Kruskal-Wallis Test, p=0.61 for moisture, p=0.16 for pH, Table 1).

**Table 1.** Soil properties in fairy circle soils.

| Position<br>in Fairy<br>Circle | Soil moisture<br>(g 100 g<sup>-1</sup>) | Organic<br>matter content<br>(%)* | Electrical<br>Conductivity<br>(EC) $\mu\text{S cm}^{-1}$* | pH |
| --- | --- | --- | --- | --- |
| Center | 0.33 ± 0.02a | 0.26 ± 0.02a | 100.8 ± 5.3a | 8.3 ± 0.08a |
| Ring | 0.32 ± 0.02a | 0.37 ± 0.02b | 171.9 ± 12.1c | 8.1 ± 0.08a |
| Matrix | 0.31 ± 0.02a | 0.34 ± 0.01b | 134.9 ± 6.6b | 8.2 ± 0.07a |

\*Indicates significant differences among fairy circle locations (ANOVA or Kruskal Wallis, n = 54 samples per position). Lower case letters indicate difference among the positions (Tukey's HSD or Dunn's Test).

### Nematode Abundance

Nematode abundance ranged from 0 - 309.2 100 g^−1^ soil. Center soils contained significantly fewer nematodes than ring or matrix soils (Kruskal Wallis Test, Dunn’s Test, p<0.0001, Fig 3A, S1 Fig). Nematode abundance was correlated positively to organic matter (p<0.001) content and EC (p=0.039) and negatively to pH (p=0.026, Spearman’s correlation,). Abundance was not correlated to moisture content (Spearman’s correlation, p>0.05).

**Fig 3.**
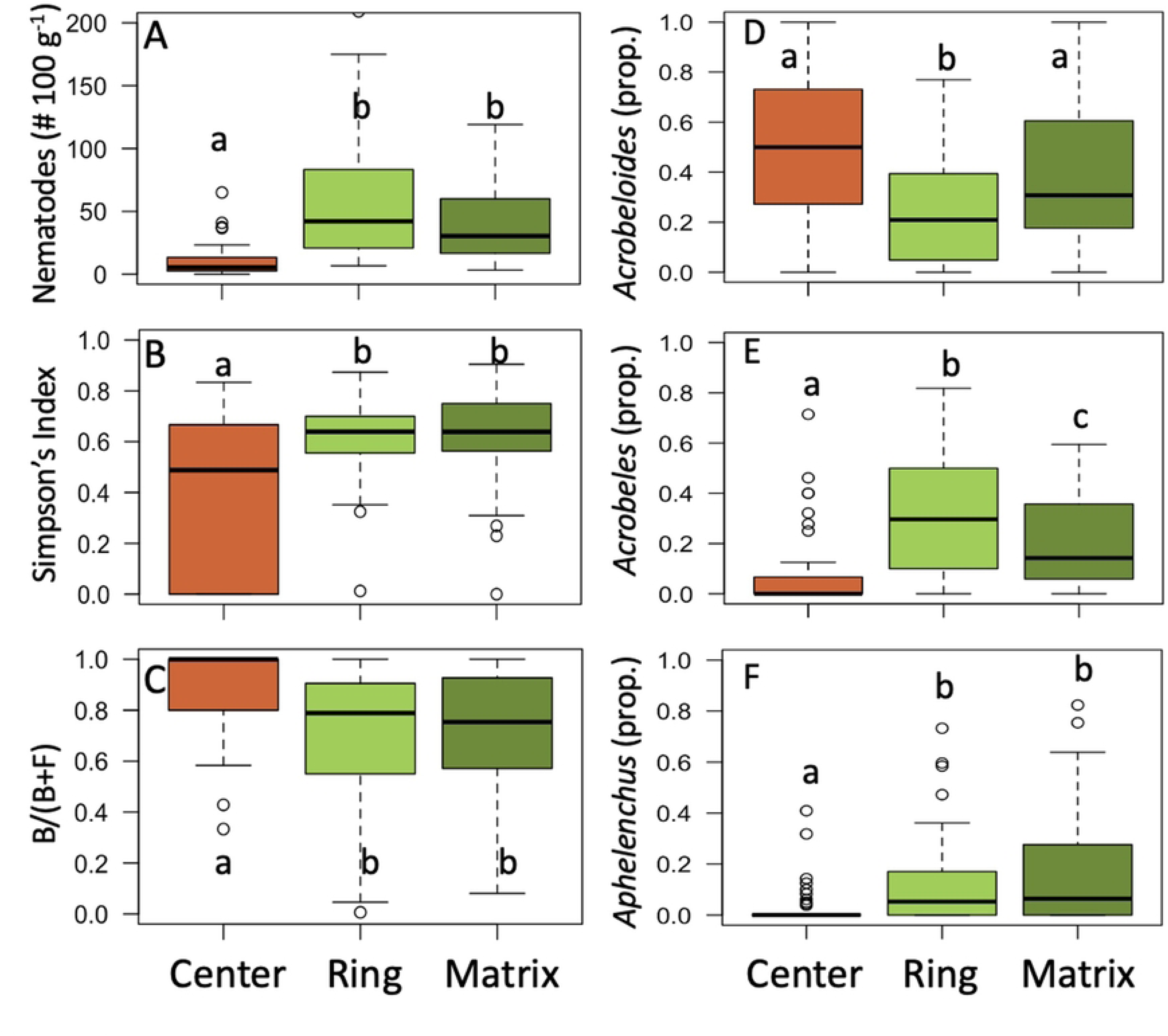
Nematode abundance and diversity in fairy circle soils collected from the center of circles, from the grass ring, or from the surrounding matrix. (A) Abundance of nematodes. (B) Simpson’s Diversity Index for nematode communities. (C) Bacterial-feeder to fungal-feeder ratio [B/(B+F)]. (D) Proportion of community: *Acrobeloides*. (E) Proportion of community: *Acrobeles*. (F) Proportion of community: *Aphelenchus*. Boxplots represent the interquartile range with lines representing the median value. Whiskers represent the minimum/maximum values, and circles are outliers. n = 54 samples per position, collected from nine field sites. Lowercase letters indicate statistically significant differences among positions (ANOVA, Tukey’s HSD Test or Kruskal-Wallis and Dunn’s Test, p<0.001).

### Nematode Diversity

Nineteen taxa of nematodes were identified across the sites (Table 2). *Acrobeles*, *Acrobeloides*, *Aphelenchus*, and *Aphelenchoides* were the only taxa present at all nine sites (Table 2). Several taxa were rare, with fewer than ten individuals seen across the study (*Carcholaimus, Discolaimus, Drilocephalobus, Nothacrobeles, Pelodera,* and *Plectus*). Richness of nematode communities ranged from 0 −10 taxa. Richness and the Simpson’s Diversity Index were both higher in ring and matrix soils than in soils from the fairy circle centers (Kruskal-Wallis Test, Dunn’s Test, *P* < 0.0001 for each, Fig 3B, S2 Fig). Richness was positively correlated to organic matter content (p<0.001) and EC (p=0.028, Spearman’s correlation), which were higher in ring and matrix soils (Table 1).

**Table 2.**
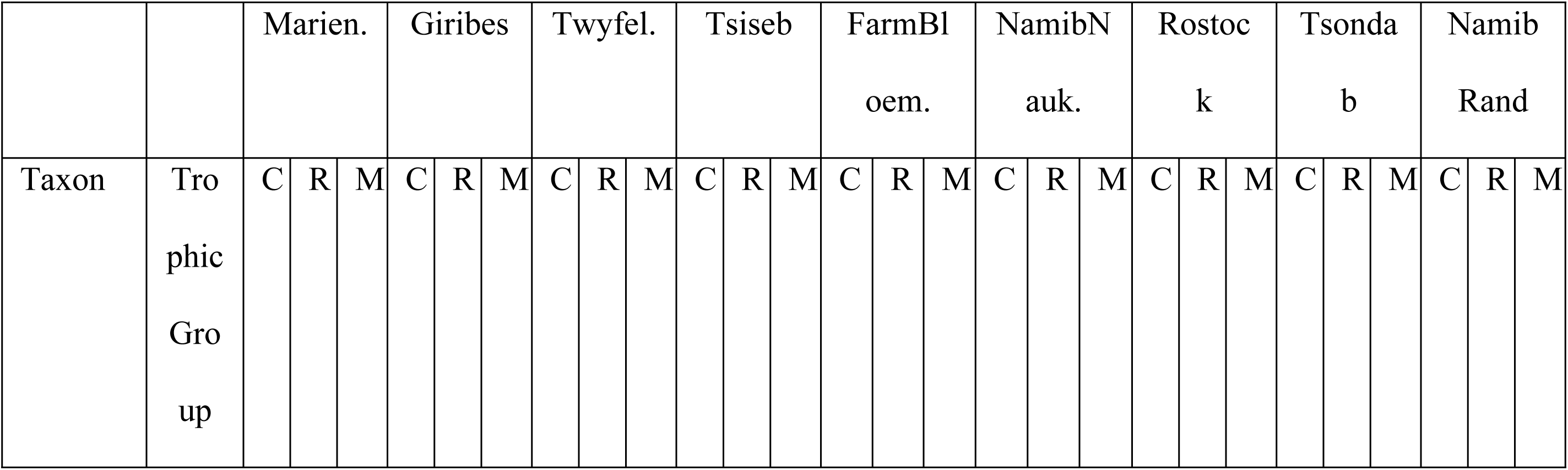
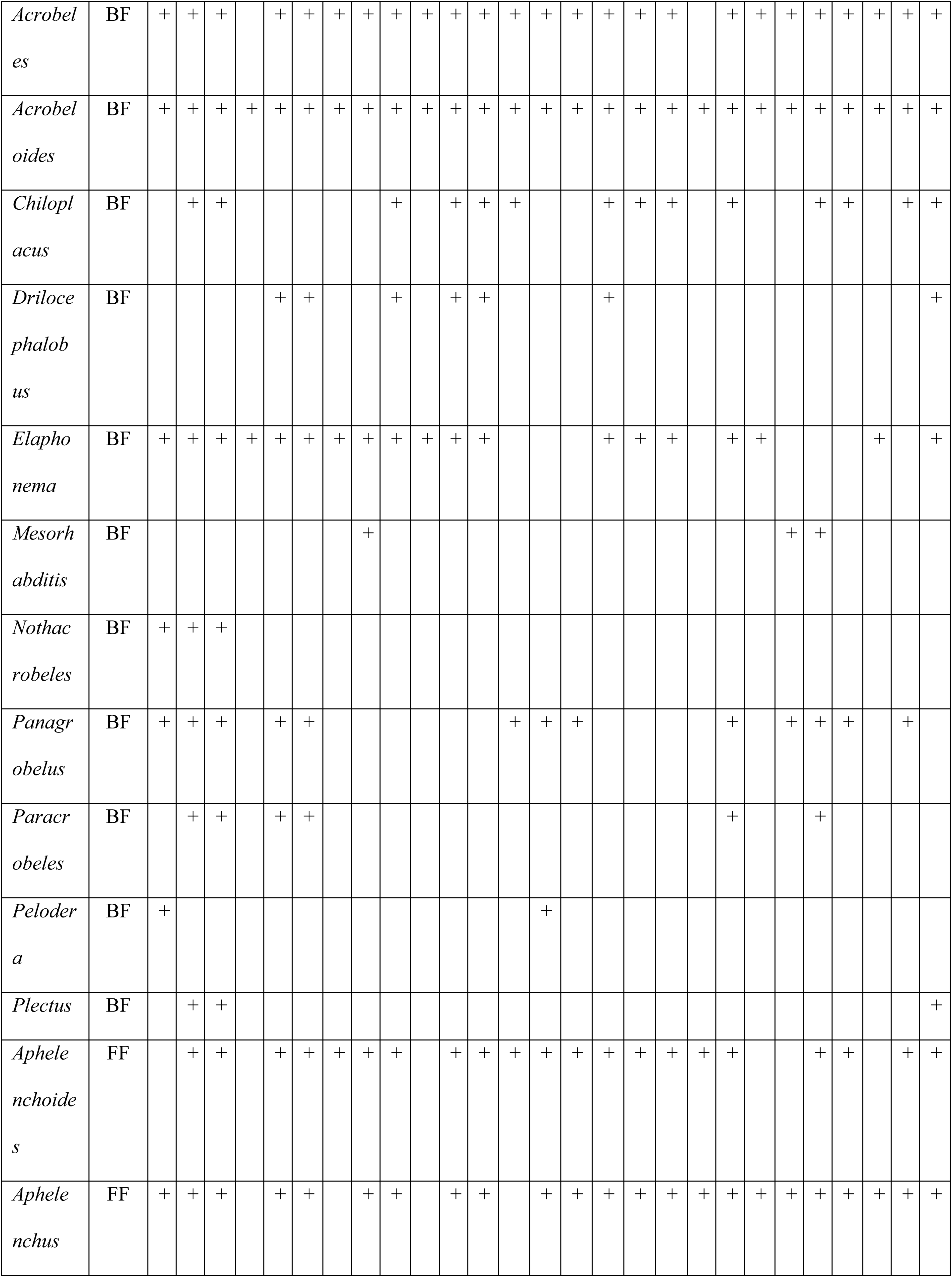
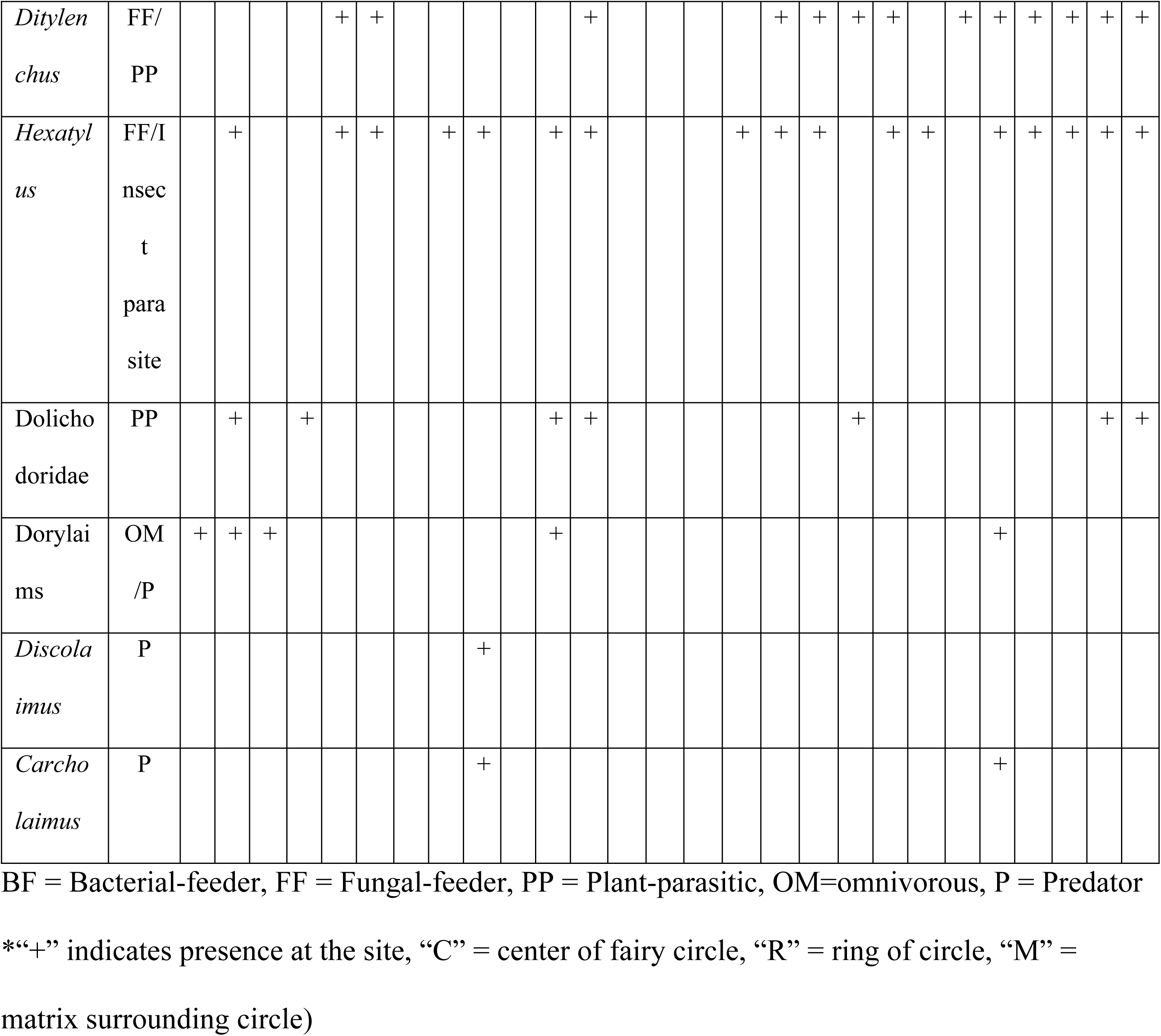
Distribution of nematode taxa among fairy circle field sites.*

### Nematode Community Structure

Nematode community structure was influenced by position in the fairy circle, at all but one of the sites (PERMANOVA, site x position interaction, p<0.001; NMDS, Fig 4A, S3 Fig). NMDS showed that center soils contained communities that were distinct from ring and matrix soils (Fig 4A). Only Farm Bloemhof lacked this separation (S3 Fig). Nematode communities in most soil samples were dominated by bacterial-feeding nematode taxa, with the proportion of the microbivore community composed of bacterial-feeders [B/(B+F)] significantly higher in the center soils than in ring or matrix (Kruskal-Wallis Test, Dunn’s Test, p<0.001, Fig 3C). *Acrobeles*, *Acrobeloides*, and *Aphelenchus* were the most common nematodes across all the fairy circle soils (i.e., 66% of nematodes identified). Their representation differed among fairy circle positions, with *Acrobeloides* over-represented and *Acrobeles* and *Aphelenchus* under-represented in center compared to ring soils (Kruskal-Wallis Test, Dunn’s Test, p<0.001 for each, Fig 3D-F).

**Fig. 4.**
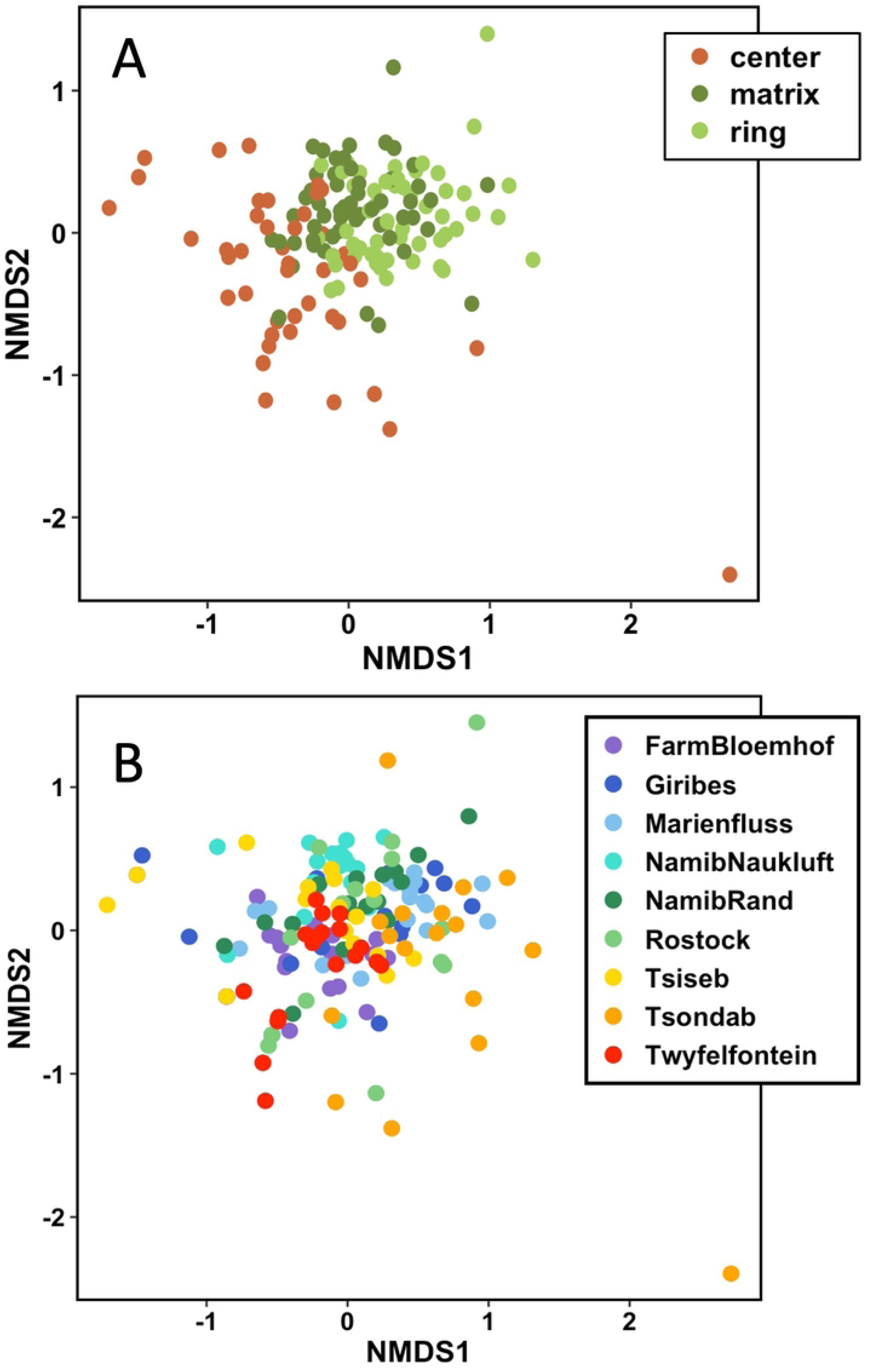
Ordination biplot from nonmetric multidimensional scaling (NMDS) analysis of soil nematode communities from fairy circles in the Namib Desert. Points represent individual samples, color coded to distinguish (A) samples collected from either the center, ring, or matrix position within fairy circles or (B) nine field sites. Stress = 0.18.

The site explained slightly more variance in nematode communities than fairy circle position (22.2% vs. 14.5%, PERMANOVA). Communities within each of the field sites clustered together to some extent, but each site also showed some overlap to several of the other sites (NMDS, Fig 4B).

RDA elucidated some of the factors contributing to differences in community composition among the positions and sites (Fig 5). The environmental variables in the model (site, position, electrical conductivity, and pH) together explained a significant proportion of the variation (26%) in community composition across sites (Permutation test, p<0.001). RDA shows that Namib Naukluft hosted more *Aphelenchus* nematodes (fungal-feeding), while Twyfelfontein, Farm Bloemhof, and the center position hosted more *Acrobeloides* (bacterial-feeders). The bacterial-feeders *Panagrobelus* and *Acrobeles* were associated with higher electrical conductivity (Fig 5). *Elaphonema* was associated with higher pH soils and with the center position (Fig 5). RDA failed to detect any connection to environmental variables for the remaining taxa, which were all rarer (Fig 5), and there remains a considerable amount of unexplained variation in the distribution of nematode taxa across these samples.

**Fig. 5.**
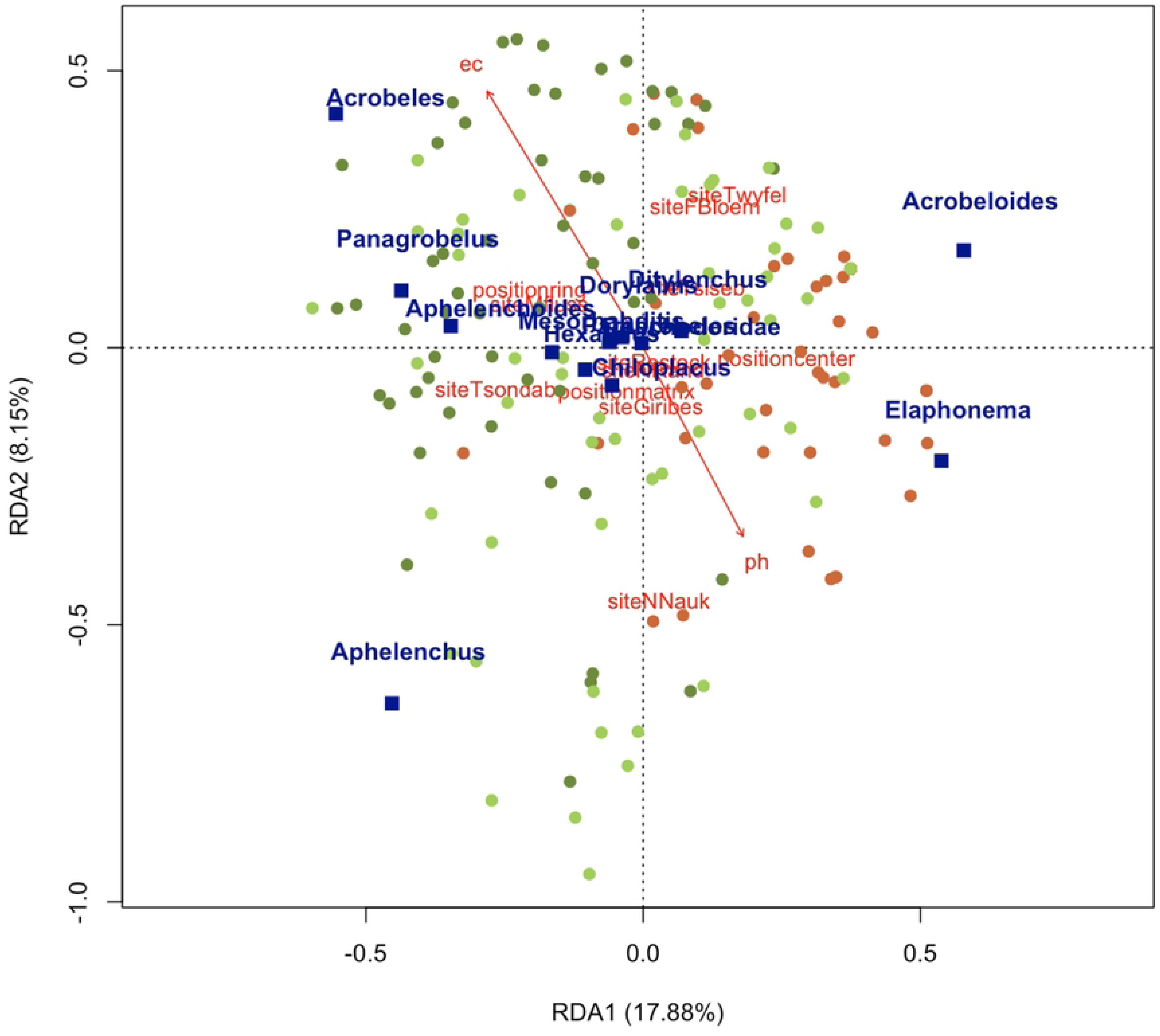
Ordination triplot for RDA of relationships among soil properties and nematode taxa in fairy circle soils. Circles represent individual soil samples for which the abundance of each taxon was determined (n=157 soils that contained nematodes). Green circles represent ring (light) and matrix (dark) soil samples, and brown circles represent samples from the centers of fairy circles. Red vectors represent the effects of explanatory variables that explained significant amounts of variation (position, site, pH, and EC). The constrained axes explained a significant proportion of the variation in nematode communities (Axis 1 explained 17.88% of the variation in the data, and Axis 2 explained 8.15%).

### Co-occurrence Network Analysis

Network analysis showed that, in general, the abundances of many of the recorded nematode taxa found were at least somewhat positively correlated with each other, suggesting that there is a core nematode community associated with fairy circle soils across the sampling sites and fairy circle positions (Fig 6A-C). This is most evident in the center soils (Fig 6A), which contained very few nematodes, but where many of those taxa co-occurred in at least some samples. Network analysis also identified taxa with stronger correlations that were most likely to co-occur. For example, dorylaims and *Nothacrobeles* were positively correlated with each other across all soil positions (Fig 6A-C).

**Fig. 6.**
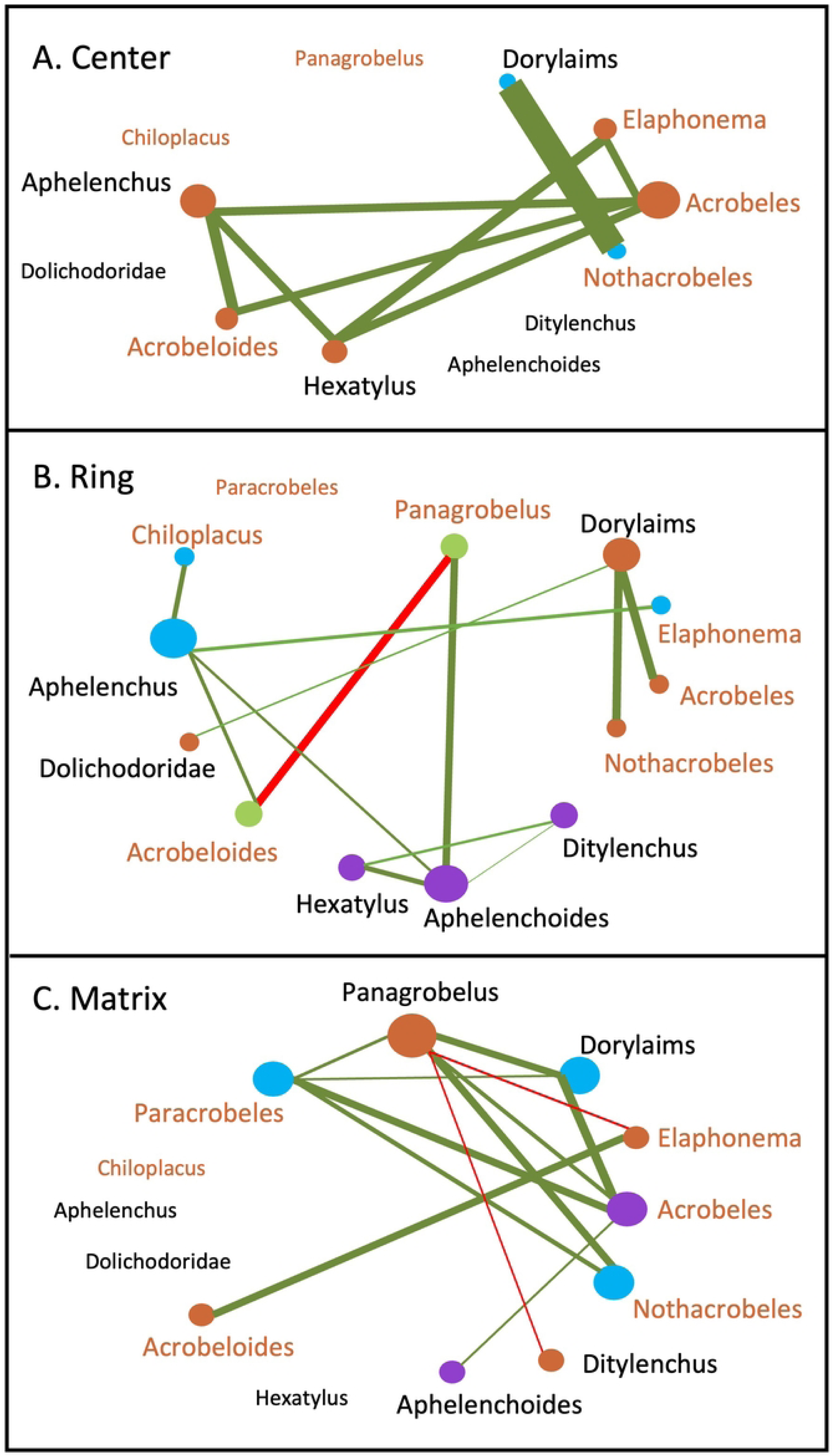
Network dynamics for nematode communities in fairy circle soils. Co-occurrence and negative correlation patterns were visualized using a threshold of 0.3 (co-occurrence) and 0.2 (negative correlations) (all p<0.05). Node size (circles) indicates the centrality of the taxon to the communities from each soil location, and node color indicates taxa that are most likely to share their higher abundances. Edges (lines) are weighted and colored according to the strength of these correlations, with negative correlations in red and positive in green. The names of bacterial-feeding taxa are in brown. (A) Center soils. (B) Ring soils. (C) Matrix soils.

In ring soils, three different assemblages of nematodes emerged (Fig 6B). Fungal-feeding *Aphelenchoides*, *Ditylenchus*, and *Hexatylus* co-occurred in many samples (Fig 6B). *Chiloplacus*, *Elaphonema*, and *Aphelenchus* also were a common assemblage. Finally, dorylaims, Dolichodoridae, *Nothacrobeles*, and *Acrobeles* were frequently found together (Fig 6B). In ring soils, there were no positive correlations among bacterial-feeding taxa (i.e., no edges, Fig 6B). Assemblages in matrix soils were less distinct than in ring soils, with several bacterial-feeding taxa showing positive correlations (Fig 6C). *Panagrobelus* was negatively correlated to two other bacterial-feeding taxa (*Elaphonema* in matrix soils and *Acrobeloides* in ring soils, Fig 6 B, C).

## Discussion

This study is the first to investigate biodiversity patterns of soil fauna across a broad geographical distribution of Namibian fairy circles and the first to examine nematode communities associated with fairy circles. Our results suggest that nematode communities respond to the unique environmental conditions created by the formation of fairy circles, but there is no evidence that nematodes are playing a role establishing or maintaining the vegetation pattern. Soils from the rings of fairy circles and the surrounding matrix support greater nematode abundance and diversity, while the centers are exceptionally depauperate. Network analysis revealed co-occurring cohorts of nematode taxa in the fairy circle environment, as well as potential niche differentiation between other taxa.

### Nematodes are connected to vegetation and organic matter

Soil organic matter content is an edaphic factor that positively affects nematodes on global and local scales [34, 38, 59]. Organic matter quality and quantity is a primary determinant of the biomass and diversity of soil bacterial and fungal communities that are consumed by many soil nematodes [60]. Prior studies of microbial communities have found that fairy circle rings and centers harbor different bacterial, archaeal, and fungal communities [16–18], with the center of the fairy circles having significantly lower microbial biomass and diversity [16].

In this study, organic matter content, which was highest in the vegetated ring and matrix soils, also was positively associated with the abundance and diversity of nematodes. In arid systems, soils from under plants have often been shown to support greater soil nematode abundance and diversity than interplant soils due to the resource island effect on soil food webs [34, 61, 62]. Although soil organic matter strongly influences nematode abundance in general, this variable was not found to explain a significant proportion of the variation in the distribution of specific taxa in our RDA analysis. This may be because all nematodes respond similarly to higher organic matter or because variation in organic content across the samples was relatively limited at the time of sampling, potentially due to the extended drought conditions. We did find that find that soils from fairy circle rings and the matrix contained more fungal-feeding and plant-parasitic taxa, demonstrating the direct role that living vegetation has on resource provisioning to nematode communities.

### Fairy circle centers are a uniquely hostile environment

We found that fairy circles have depauperate centers with extremely low soil nematode abundance and diversity, averaging just 9 nematodes 100 g^−1^ soil across nine field sites (range 0 - 65, n = 54). The paucity of nematodes in the center soils is striking compared to studies of nematode communities in other unvegetated soils in the region. For example, across the gravel plains in the hyperarid Namib Desert, bare soils in between shrubs harbored an average of 57.7 ± 25.6 nematodes 100 g^−1^ soil (range 0 - 525, n = 27, [48]). A similar result was found for interplant spaces between *Welwitschia* plants in the Namib, where nematode density averaged 34.7 ± 6.4 nematodes 100 g^−1^ soil (range 0.8 - 198, n =36, [63]). In both of those studies, differences between soil nematode abundance under plants and in interplant spaces were not discernable at many sites [48, 63].

In the Namib Desert gravel plains, the sporadic germination and growth of desert ephemerals in response to significant rainfall events seems to help to support nematode abundance in areas between perennial plants. Interplant spaces in the Namib Desert that are closer to the Atlantic coast also harbor fog-moistened lichens, cryptobiotic crusts, and endoliths [48, 64–66]. These photosynthetic organisms appear to be sufficiently productive to support soil food webs in interplant spaces, as has been found in other arid ecosystems [67, 68]. In contrast, the mechanisms that prevent vascular plants from growing in fairy circle centers also may prohibit the development of cryptobiotic crusts, ultimately resulting in a nearly inhabitable zone for nematodes. Ripples typically are visible in the sand in the centers of fairy circles (Fig 1A), suggesting that there is quite a bit of wind disturbance of the surface soils, which would make crust formation challenging. Center soils also have been found to contain more coarse particles than those found in the rings, due to wind activity [28, 37]. In a study of dune sands in the Namib, non-vegetated soils also contained very few nematodes, supporting the possible impact of wind disturbance on habitability [61].

The small number of nematodes that were found in center soils might be part of established populations that feed and reproduce there, rather than transients that were blown in from ring or matrix soils. We found that *Acrobeloides*, a bacterial-feeder, is over-represented in the center soil communities compared to the ring soils. *Acrobeloides* may be suited to live the hostile center environment compared to other nematode taxa, although the low numbers suggest the environment remains challenging. *Acrobeloides* are generally described as being bacterial-feeding, but some species have been found in association with insects [69–71]. A possible interaction of this nematode with the termites or ants that are often observed to be active in fairy circles warrants future investigation. Microbivorous nematodes, including *Acrobeloides*, have been found associated with termite bodies, although their impact on the termites is not well-characterized [72, 73]. Alternatively, the rings and centers of fairy circles harbor distinct phylotypes of bacteria [18], and *Acrobeloides* may be adapted to consume bacteria found in center soils. All these factors may collectively contribute to *Acrobeloides* relative success in fairy circle centers.

### Extremely low moisture and high temperatures have minimal direct impacts on nematodes in fairy circle soils

In this study, soil moisture was exceptionally low at all sites due to drought conditions (∼ 0.3%), and no differences were detected between center, ring, or matrix soils. Although not measured in this study, prior work has shown that fairy circle centers can have higher soil moisture due to the lack of plant uptake [2]. Nematodes rely on water films for movement, feeding, and reproduction [74], but under dry conditions, they can enter into an inactive and ametabolic state known as anhydrobiosis [75]. Soil nematodes in desert ecosystems can rapidly become active in response to rainfall [76], melting snowfall [77], and fog that wets the soil [48]. The absence of moisture for long periods does not appear to be a factor directly affecting nematode abundance and diversity in fairy circle soils, although nematodes are likely to be affected indirectly due to reduced plant productivity [48].

Soil temperatures can be quite high in fairy circle soils and have been shown to vary among positions. In a study at Namib Rand in February (austral summer), the surface of center soils had an average daytime temperature of 45°C while the matrix averaged 47°C, and daily maximum temperatures exceeded 55°C [36]. Soils were somewhat cooler at 15 cm depth (i.e., maximum < 50°C [36]). Nematodes in anhydrobiosis are temperature resistant, with most showing survival at 50°C for one hour in a laboratory experiment (longer exposures were not tested [78]). It does not seem likely soil temperature is contributing to the hostile environment of the fairy circle centers, as they have been found to be cooler than vegetated soils. Overall, the abundance and diversity nematodes in fairy circle soils seems to be affected more by differences in the distribution of plants and plant-derived organic matter than by differences in soil moisture or temperature.

### Insights from network analysis of nematode communities in fairy circle soils

Belowground communities are exceptionally diverse, and nematodes are the most abundant animal on the planet [38, 79]. Untangling the factors that influence which taxa are present in communities is therefore a challenging task. Community assembly patterns in nematodes have not been explored often, but deterministic processes based on abiotic and biotic interactions have been shown to be important [46, 79]. Co-occurrence network analysis is an emerging tool that has been used to understand nematode communities [80–85].

Our network analysis of nematode assemblages in fairy circle soils found that most nematode taxa that exist in these soils can occur together. However, there was variation in taxa abundance among samples, allowing us to identify assemblages of nematode most likely to co-occur and thrive in the same environment. Across all fairy circle positions, soils with higher abundances of dorylaims were more likely to have more *Nothacrobeles,* suggesting that they have overlapping ecological niches. Species of dorylaims are usually omnivorous or predatory, and *Nothacrobeles* are bacterial-feeding [30], so niche overlap is not due to a shared food source, although it’s possible *Nothacrobeles* are dorylaim prey. Both nematodes were rare across the sites, but they specifically were found in many of the samples collected from the Marienfluss site, suggesting the role of environmental factors in influencing their co-occurrence. This site had the lowest mean pH of all the sites (7.3 vs. 7.6 - 8.8 for the other sites).

In ring soils, fungal-feeding *Aphelenchoides*, *Ditylenchus*, and *Hexatylus* were positively correlated, likely due to enhanced fungal biomass in these plant-influenced soils. In ring soils, there were no positive correlations among bacterial-feeding nematode taxa, suggesting that their distribution here could be stochastic. This is a possible outcome of the taller ring grasses capturing more wind-blown nematodes [86]. There were some positive correlations among bacterial-feeding taxa in matrix soils, where the grass typically does not grow as tall, and where deterministic processes may have larger influence.

*Panagrobelus* demonstrated niche specialization by having negative correlations with two other bacterial-feeding taxa (*Elaphonema* in matrix soils and *Acrobeloides* in ring soils). RDA shows that *Panagrobelus* are more abundant in soils with higher electrical conductivity (i.e., salinity) than *Acrobeloides* or *Elaphonema*, suggesting that adaptations to edaphic factors may be contributing to this specialization. Biotic interactions, such as competition for a shared food source, may also be contributing, however. Caruso et al. [45] studied nematode communities consisting of up to three species in the soils of the Antarctic Dry Valleys. They found that the presence of one bacterial-feeding species in a soil (*Plectus*) decreased the probably of the other bacterial-feeder (*Scottnema*) being present. *Scottnema* was found primarily in the driest, lowest organic matter, and high salinity soils, and *Plectus* occurs in soils with higher moisture [45, 67]. While *Scottnema* could survive in *Plectus*’ less extreme habitat, it was less likely to be found there, and if present, less abundant, possibly due to being less competitive for food in this environment [45]. It was suggested that *Plectus* may not have the ability to survive as well in the dry, salty soils where *Scottnema* flourishes [45]. In Namib Desert soils, *Elaphonema* is a fantastical nematode with elaborate, “elephantine” mouth parts [87]. Although this nematode was present at seven of the nine field sites, it was most abundant in the Namib Naukluft samples. This site was distinct from the other sites due to its westernmost location in the gravel plains of the Namib Desert, where conditions are generally drier and the soils contain more gravel than at sandier sites further inland. *Acrobeloides*, on the other hand, was widely distributed and abundant across the sites, except in soils with many *Panagrobelus*, suggesting competition may limit their co-occurrence. Similar to *Plectus*/*Scottnema* in Antarctica, these three bacterial-feeders in the Namib Desert seem to be showing habitat specialization as a function of biotic and abiotic interactions.

Network analysis, combined with RDA, suggests that differences in soil physico-chemical properties as well as biotic interactions influence the distribution of different taxa in fairy circle soils, resulting in the formation of common assemblages. Information on the specific ecologies of nematode taxa is relatively limited. The results of network analysis can guide future research to understand the mechanisms underlying nematode species coexistence or exclusion, which may include physiological or life history adaptations to specific environments and/or resource niche differentiation.

### Future opportunities for research

The current study represents a snapshot of diversity in a specific layer of the soil. As such, we did not detect significant numbers of nematode plant-parasites, but our sampling regime did not focus on roots. We focused on the uppermost layer of soil because we wanted to study the environment where plants germinate and die, resulting in the bare centers of fairy circles [88], and where significant grass litter decomposition occurs [89–90]. Nematodes are likely to be present in deeper layers of the soil, representing an opportunity for future study. We did conduct some deeper sampling at one site (10-20 cm, Farm Bloemhof), where we found similar abundances of nematodes to the upper layer in center soils, but with higher numbers in the deeper layers of ring soils (mainly more *Aphelenchus,* a fungal-feeder). Deeper soil sampling and/or direct sampling of roots for extraction of nematodes may elucidate a broader diversity of nematodes, including more plant ectoparasites.

Termite or ant activity has been posited as one cause for the formation of fairy circles [10], and we did observe evidence of both insects at many of the sites, even though we did not specifically record their presence. We did not detect any nematode taxa that are known specifically to be termite parasites. Species of *Hexatylus* are described as either fungal-feeders and/or entomopathogens [53], but we could not find any reports of their presence in termites or ants. Many nematodes show associations with insect bodies [91]; as noted above for *Acrobeloides*) and are known to engage in phoresy or to consume bacteria within corpses. A study of the nematodes associated with arthropods at fairy circles may reveal a new layer of complexity to the ecological interactions underpinning fairy circle patterning.

## Conclusions

The mechanisms behind the creation and maintenance of fairy circles remain enigmatic, but this study demonstrates that the vegetation patterning has belowground impacts. Soil nematode communities show distinct configurations that are connected to the fairy circle vegetation, primarily due to the impact of plants on soil organic matter, pH, and salinity. Although non-vegetated Namib Desert soils have been shown by prior work to consistently harbor robust nematode communities in other locations, fairy circle soils are nearly defaunated, suggesting that same mechanism that limits establishment of plants in the circle also is affecting soil nematodes, either directly or indirectly.

## Acknowledgements

We thank the Namibia University of Science and Technology for logistical support, especially Elizabeth Nashidengo and Shapopi Kamanja. We also warmly thank the proprietors and staff at Africa on Wheels, Solitaire Roadhouse, Tsondab Valley Reserve, Rostock Ritz, Namib Rand Nature Reserve, and Farm Bloemhof.

## Supporting information

**S1 Fig.**
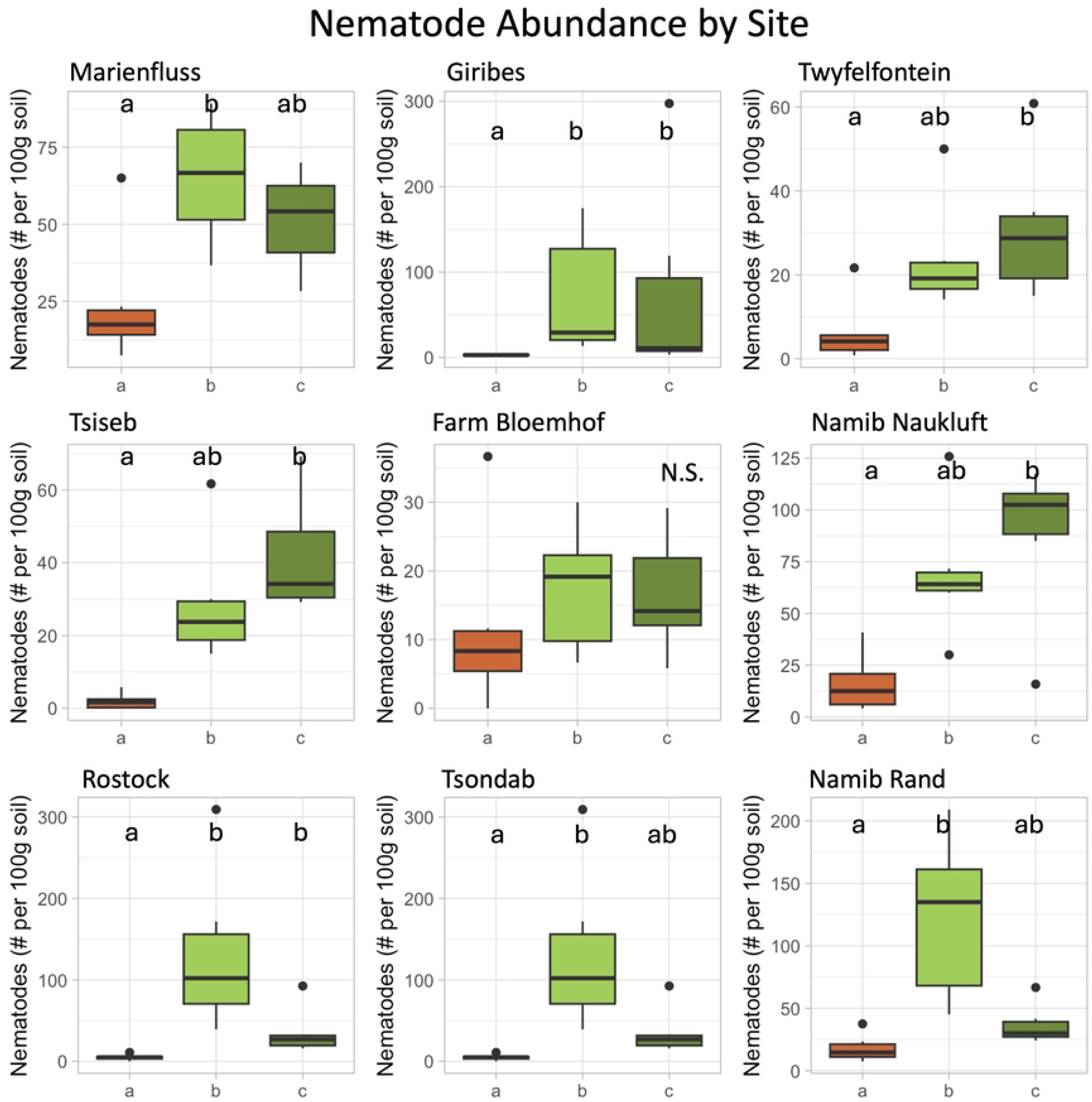
Nematode abundance in fairy circle soils from nine field sites. Boxplots represent the interquartile range with lines representing the median value. Whiskers represent the minimum/maximum values, and circles are outliers. n = 6 samples per position. Lowercase letters indicate statistically significant differences among positions (N.S. = not significant, Kruskal-Wallis and Dunn’s Test, *P* < 0.05). a= center, b = ring, and c = matrix soils.

**S2 Fig.**
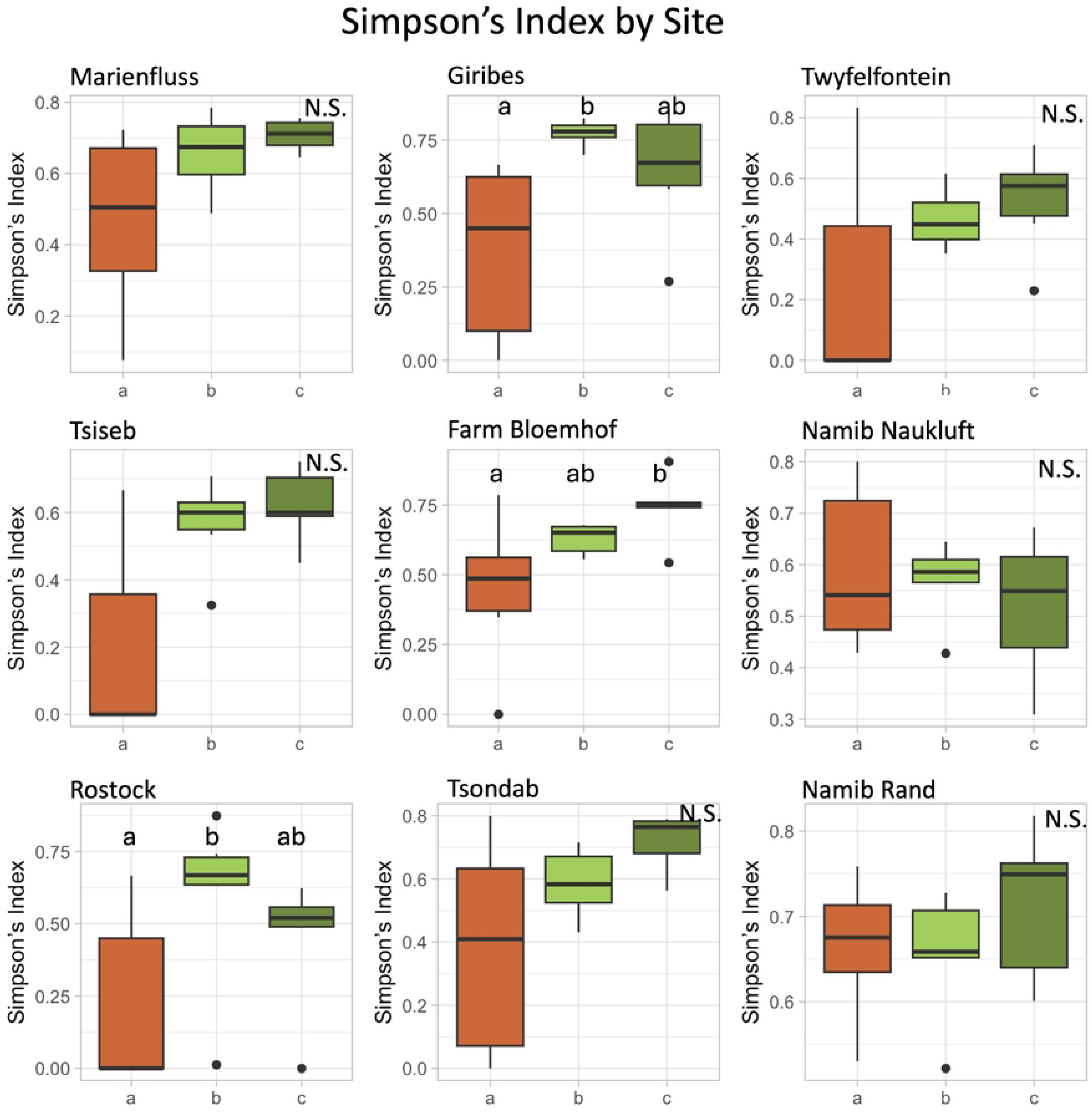
Simpson’s Index of diversity for nematode communities in fairy circle soils from nine field sites. Boxplots represent the interquartile range with lines representing the median value. Whiskers represent the minimum/maximum values, and circles are outliers. n = 6 samples per position. Lowercase letters indicate statistically significant differences among positions (N.S. = not significant, Kruskal-Wallis and Dunn’s Test, *P* < 0.05). a= center, b = ring, and c = matrix soils.

**S3 Fig.**
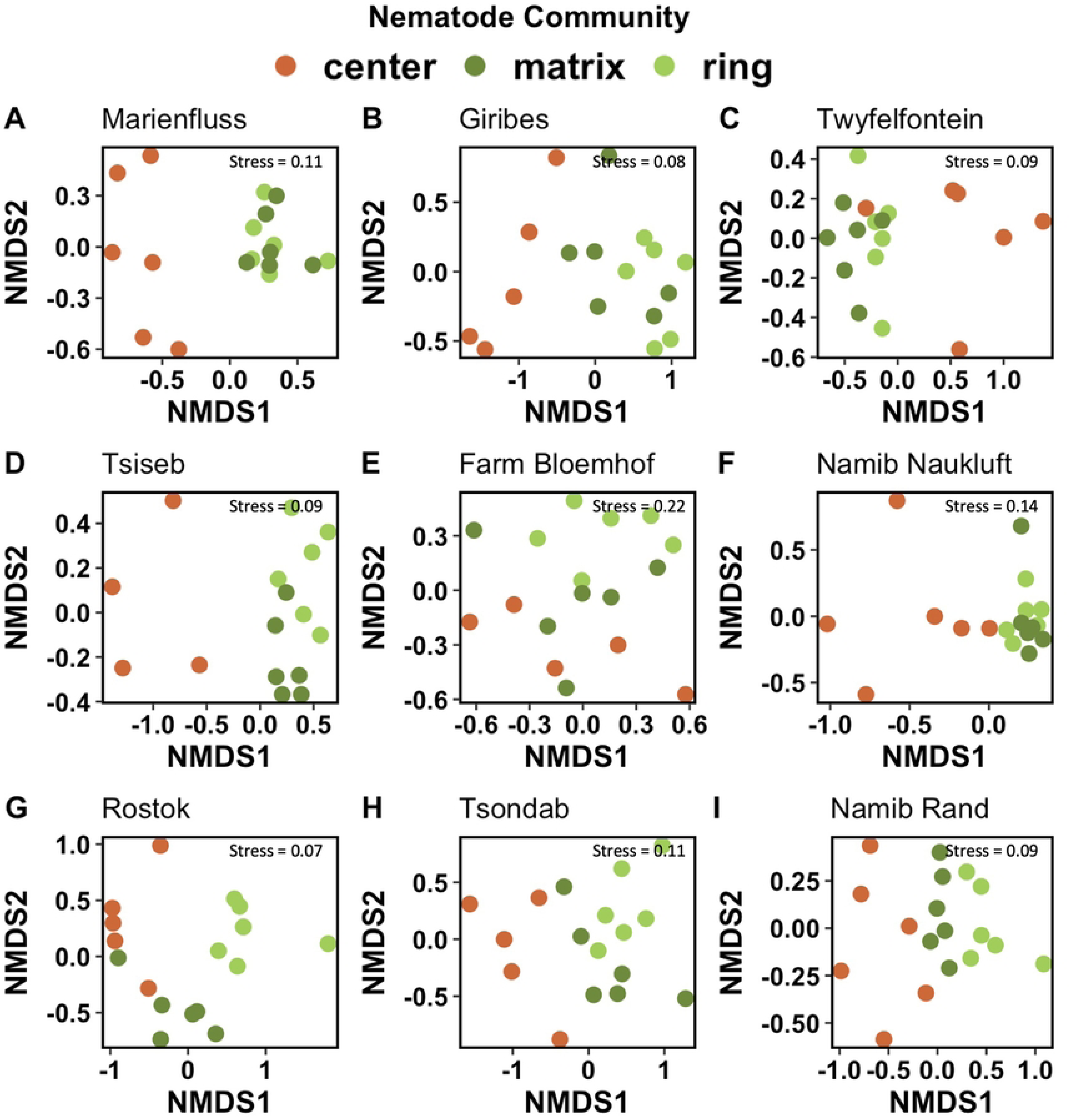
NMDS of Bray-Curtis Dissimilarity for soil nematode communities in fairy circle soils from nine field sites. Center = brown dots, ring = light green dots), matrix = dark green dots.

## References

1. Tinley KL. Etosha and the Kaokoveld. Afr Wild Life. 1971;25:1–16.

2. Jürgens, N. The biological underpinnings of Namib Desert fairy circles. Nature. 2013;339:1618–21.

3. Getzin S, Yizhaq H, Tschinkel W. Definition of “fairy circles” and how they differ from other common vegetation gaps and plant rings. J Veg Sc. 2021;32:e13092.

4. Meyer JJM, Schutte CS, Galt N, Hurter JW, Meyer NL. The fairy circles (circular barren patches) of the Namib Desert - What do we know about their cause 50 years after their first description? S Afr J Bot. 2021;140:1–14.

5. Getzin S, Yizhaq H, Bell B, Erickson T, Postle A, Katra I, et al. Discovery of fairy circles in Australia supports the self-organisation theory. Proc Natl Acad Sci USA. 2016;113:3551–6.

6. Guirado E, Delgado-Baquerizo M, Benito BM, Molina-Pardo JL, Berdugo, M, Martínez-Valderrama JM, et al. The global biogeography and environmental drivers of fairy circles. Proc Natl Acad Sci USA. 2023;120:e2304032120.

7. van Rooyen MW, Theron GK, van Rooyen N, Jankowitz WJ, Matthews WS. Mysterious circles in the Namib Desert: Review of hypotheses on their origin. J Arid Environ. 2004;57:467–85.

8. Topographic basemap. Scale Not Given. “World Topographic Map” [Internet]. Redlands (CA): Esri. c2020 – [cited 2024 November 1]. Available from: http://www.arcgis.com/home/item.html?id=30e5fe3149c34df1ba922e6f5bbf808f

9. Tarnita CE, Bonachela JA, Sheffer E, Guyton JA, Coverdale TC, Long RA, et al. A theoretical foundation for multi-scale regular vegetation patterns. Nature. 2017; 541:398–401.

10. Jüergens N, Groengroeft A, Gunter F. Evolution at the arid extreme: The influence of climate on sand termite colonies and fairy circles of the Namib Desert. Philos T Roy Soc B. 2023;378:20220149.

11. Getzin S, Holch S, Ottenbreit J, Yizhaq H, Wiegand K. Spatio-temporal dynamics of fairy circles in Namibia are driven by rainfall and soil infiltrability. Landscape Ecol. 2024;39:122.

12. Albrecht CF, Joubert JJ, DeRycke PH. Origin of the enigmatic, circular, barren patches (‘Fairy Rings’) of the pro-Namib. S Afr J Sci. 2001;97:23–27.

13. Naudé Y, van Rooyen MW, Rohwer ER. Evidence for a geochemical origin of the mysterious circles in the Pro-Namib desert. J Arid Environ. 2011;75:446–56.

14. Theron GK. Die verskynsel van kaal kolle in Kaokoland, SuidWes-Afrika. J S Afr Biol Soc. 1979;20:43–53. Afrikaans.

15. Meyer JM, Schutte CE, Hurter JW, Galt NS, Degashu P, Breetzke G, et al. The allelopathic, adhesive, hydrophobic and toxic latex of *Euphorbia* species is the cause of fairy circles investigated at several locations in Namibia. BMC Ecol. 2020;20:45.

16. Eicker A, Theron GK, Grobbelaar N. ’n Mikrobiologiese studie van ‘kaal kolle’ in die Giribesvlakte van Kaokoland, S.W.A.-Namibië. S Afr J Bot. 1982;1:69–74. Afrikaans.

17. Ramond J-B, Pienaar A, Armstrong A, Seely M, Cowan DA. Niche-partitioning of edaphic microbial communities in the Namib Desert gravel plain fairy circles. PLOS One. 2014;9:e109539.

18. van der Walt AJ, Johnson RM, Cowan DA, Seely M, Ramond J-B. Unique microbial phylotypes in Namib desert dune and gravel plain fairy circle soils. Appl Environ Microbiol. 2016;82:4592–601.

19. Cramer MD, Barger NN. Are Namibian “Fairy Circles” the consequence of self-organizing spatial vegetation patterning? PLOS One. 2013;8:e70876.

20. Fernandez-Oto C, Tlidi M, Escaff D, Clerc MG. Strong interaction between plants induces circular barren patches: fairy circles. Philos T Roy Soc A. 2014;372:20140009.

21. Getzin S, Wiegand K, Wiegand T, Yizhaq H, von Hardenberg J, Meron E. Adopting a spatially explicit perspective to study the mysterious fairy circles of Namibia. Ecography. 2015;38:1–11.

22. Getzin S, Holch S, Yizhaq H, Wiegand K. Plant water stress, not termite herbivory, causes Namibia’s fairy circles. Perspect Plant Ecol. 2022;57:125698.

23. Moll EJ. The origin and distribution of fairy rings in Namibia. In: Seyani JH, Chikuni AC, editors. Proceedings of the 13th Plenary Meeting AETFAT, Malawi. 2nd ed. Zomba: National Herbarium and Botanic Gardens of Malawi; 1994. p. 1203–09.

24. Picker MD, Ross-Gillespie V, Vlieghe K, Moll E. Ants and the enigmatic Namibian fairy circles – cause and effect? Ecol Entom. 2012;37:33–42.

25. Jüergens N. Exploring common ground for different hypotheses on Namib fairy circles Ecography. 2015;38:12–14.

26. Vlieghe K, Picker M, Ross-Gillespie V, Erni, B. Herbivory by subterranean termite colonies and the development of fairy circles in SW Namibia. Ecol Entom. 2015;40:42–49.

27. Jüergens N, Gröngröft A. Sand termite herbivory causes Namibiá s fairy circles – A response to Getzin et al. (2022). Perspect Plant Ecol. 2023;60:125745.

28. Ravi S, Wang L, Kaseke KF, Buynevich IV, Marais E. Ecohydrological interactions within “fairy circles” in the Namib Desert: Revisiting the self-organization hypothesis, J Geophys Res-Biogeo. 2017;122:405–14.

29. Jiang Y, Qian H, Wang X, Chen L, Liu M, Li H, et al. Nematodes and microbial community affect the sizes and turnover rates of organic carbon pools in soil aggregates. Soil Biol Biochem. 2018;119:22–31.

30. Yeates GW, Bongers, T, De Goede RG, Freckman DW, Georgieva SS. Feeding habits in soil nematode families and genera-an outline for soil ecologists. J Nematol. 1993;25:315–31.

31. Wardle DA, Bardgett RD, Klironomos JN, Setälä H, van der Putten WH, Wall DH. Ecological linkages between aboveground and belowground biota. Science. 2004;304: 1629–33.

32. Tsiafouli MA, Thébault E, Sgardelis SP, de Ruiter PC, van der Putten WH, Birkhofer K, et al. Intensive agriculture reduces soil biodiversity across Europe. Glob Change Biol. 2014;21:973–85.

33. Porazinska DL, Bueno de Mesquita CP, Farrer EC, Spasojevic MJ, Suding KN, Schmidt SK. Nematode community diversity and function across an alpine landscape undergoing plant colonization of previously unvegetated soils. Soil Biol Biochem. 2021;161:108380.

34. Treonis AM, Sutton KA, Unangst SK, Wren JE, Dragan ES, McQueen JP. Soil organic matter determines the distribution and abundance of nematodes on alluvial fans in Death Valley, CA. Ecosphere. 2019;10:e02659.

35. Cramer M, Barger N, Tschinkel R. Edaphic properties enable facilitative and competitive interactions resulting in fairy circle formation. Ecography. 2017;40:1210–20.

36. Vlieghe K, Picker M. Do high soil temperatures on Namibian fairy circle discs explain the absence of vegetation? PLOS One. 2019;14:e0217153.

37. Yizhaq H, Rein C, Saban L, Cohen N, Kroy K, Katra I. Aeolian sand sorting and soil moisture in arid Namibian fairy circles. Land. 2024;13:197.

38. van den Hoogen J, Geisen S, Routh D, Ferris H, Traunspurger W, Wardle DA. Soil nematode abundance and functional group composition at a global scale. Nature. 2019;572:194–98.

39. Bongers T. The Maturity Index, the evolution of nematode life history traits, adaptive radiation and cp-scaling. Plant Soil. 1999;212:13–22.

40. Ferris H, Bongers T, De Goede RGM. A framework for soil food web diagnostics: Extension of the nematode faunal analysis concept. Appl Soil Ecol. 2001;18:13–29.

41. De Deyn GB, Raaijmakers CE, Ruijven J, Berendse F, Putten VD. Plant species identity and diversity effects on different trophic levels of nematodes in the soil food web. Oikos. 2004;106:576–86.

42. Liang W, Lou Y, Li Q, Zhong S, Zhang X, Wang J. 2009. Nematode faunal response to long-term application of nitrogen fertilizer and organic manure in Northeast China. Soil Biol Biochem. 2009;41:883–90.

43. Treonis AM, Unangst SK, Kepler RM, Buyer JS, Cavigelli MA, Mirsky SB, Maul JE. Characterization of soil nematode communities in three cropping systems through morphological and DNA metabarcoding approaches. Sci Rep-UK. 2018;8:2004.

44. Thakur MP, Phillips HRP, Brose U, De Vries FT, Lavelle P, Loreau M, et al. Towards an integrative understanding of soil biodiversity. Biol Rev Camb Philos. 2020;95:350–64.

45. Caruso T, Hogg ID, Nielsen UN, Bottos EM, Lee CK, Hopkins DW, et al. Nematodes in a polar desert reveal the relative role of biotic interactions in the coexistence of soil animals. Commun Biol. 2019;2:63.

46. Gattoni K, Gendron, EMS, McQueen, JP, Powers, K, Powers, TO, Harner, MJ. The nature of microbial diversity and assembly in the Nebraska Sandhills depends on organismal identity and habitat type. Front Ecol Evol. 2024;12:1305930.

47. Freckman, DW, Virginia RA. Low-diversity Antarctic soil nematode communities: Distribution and response to disturbance. Ecology. 1997;78:363–69.

48. Treonis AM, Marais E, Maggs-Kölling G. Nematode communities indicate diverse soil functioning across a fog gradient in the Namib Desert gravel plains. Ecol Evol. 2022;12:e9013.

49. Gardner WH. Water content. In: Klute A, editor. Methods of soil analysis, Part 1. Physical and mineralogical methods. Madison, WI: American Society of Agronomy, Soil Science Society of America; 1986. p. 493–544.

50. Nelson DW, Sommers LE. Total carbon, organic carbon, and organic matter. In: Sparks DL, Page AL, Helmke PA, et al., editors. Methods of soil analysis, Part 3, Chemical methods. Madison, WI: American Society of Agronomy, Soil Science Society of America; 1996. p. 961–1010.

51. Ingham RE. Nematodes. In: Weaver, et al., editors. Methods of soil analysis, Part 2, Microbiological and biochemical properties. Madison, WI: American Society of Agronomy. Soil Science Society of America; 1994. p. 459–90.

52. Bongers T. De Nematoden van Nederland. Utrecht: Stichting Uitgeverij Koninklijke Nederlandse Natuurhistorische Vereniging. 1988.

53. Andrássy I. Free-living nematodes of Hungary I. (*Nematoda errantia*.) In: Csuzdi, CS, Mahunka, S, editors. Pedozoologica Hungarica, No. 3, Budapest, Hungarian Natural History Museum. 2005.

54. Simpson EH. Measurement of diversity. Nature. 1949;163:688.

55. R Development Core Team. R: A language and environment for statistical computing. Version 4.4 [software]. 2024. Available from: https://www.R-project.org/.

56. Oksanen J, Guillaume Blanchet F, Friendly M, Kindt R, Legendre P, McGlinn D, et al. vegan: Community ecology package. 2018. Available from: https://cran.r-project.org/web/packages/vegan/index.html

57. Rao CR. The use and interpretation of principal component analysis in applied research. Sankhya Ser A. 1964;26:329–358.

58. Newman MEJ. Assortative mixing in networks. Phys Rev Lett. 2002;89:208701.

59. Yeates GW. Effects of plants on nematode community structure. Annu Rev Phytopathol.1999;37:127–49.

60. Fierer N. Embracing the unknown: Disentangling the complexities of the soil microbiome. Nat Rev Microbiol. 2017;15:579–90.

61. Marais E, Maggs-Kölling G, Sherman C, Doniger T, Liu R, Tripathi BM, et al. Profiling soil free-living nematodes in the Namib Desert, Namibia. J Arid Land 2020;12:130–43.

62. Gan HY, Hohberg K, Schneider, C, Ebner M, Marais E, Miranda T, et al. The hidden oases: Unveiling trophic dynamics in Namib’s fog plant ecosystem. Sci Rep-UK. 2024;14;13334.

63. Treonis AM, Marais E, Maggs-Kölling G. Soil nematode communities vary among populations of the iconic desert plant, *Welwitschia mirabilis*. Pedobiologia. 2024;103: 150943.

64. Qu EB, Omelon CR, Oren A, Meslier V, Cowan DA, Maggs-Kölling G, et al. Trophic selective pressures organize the composition of endolithic microbial communities from global deserts. Front Microbiol. 2020;10:2952.

65. de los Ríos A, Garrido-Benavent I, Limón, A, et al. Novel lichen-dominated hypolithic communities in the Namib Desert. Microb Ecol. 2022;83:1036–48.

66. Bosch J, Lebre PH, Marais E, Maggs-Kölling G, Cowan DA. Kinetics and pathways of sub-lithic microbial community (hypolithon) development. Environ Microbiol Rep. 2024;16:e13290.

67. Treonis AM, Wall DH, Virginia RA. Invertebrate biodiversity in Antarctic Dry Valley soils and sediments. Ecosystems. 1999;2:482–92.

68. Darby BJ, Neher DA, Belnap J. Soil nematode communities are ecologically more mature beneath late-than early-successional stage biological soil crusts. Appl Soil Ecol. 2007;35:203–212.

69. Azizoglu U, Karabörklü S, Ayvaz A, Yilmaz S. Phylogenetic relationships of insect– associated free–living rhabditid nematodes from eastern mediterranean region of Turkey. Appl Ecol Env Res. 2016;14:93–103.

70. Rana A, Bhat AH, Bhargava S, Chaubey AK, Abolafia J. Morphological and molecular characterization of *Acrobeloides saeedi* Siddiqi, De Ley and Khan, 1992 (Rhabditida, Cephalobidae) from India and comments on its status. J Nematol. 2020;52:1–21.

71. Bhat AH, Loulou A, Abolafia J, Machado RAR, Kallel S. Comparative morphological and molecular analyses of *Acrobeloides bodenheimeri* and *A. tricornis* Cobb, 1924 (Rhabditida, Cephalobidae) from Tunisia. Nematology 2023;25: 207–26.

72. Wang C, Powell JE, OConnor BM. Mites and nematodes associated with three subterranean termite species (Isoptera: Rhinotermitidae). Fla Entomol. 2002;85:499–506.

73. Carta LK, Handoo ZA, Lebedeva NI, Raina AK, Zhuginisov TI, Khamraev AS. *Pelodera termitis sp. nov*. and two other rhabditid nematode species associated with the Turkestan termite *Anacanthotermes turkestanicus* from Uzbekistan. Int J Nematol. 2010;20:125–34.

74. Yeates GW. Ecological behavioural adaptations. In: Gaugler R, Bigrami AL, editors. Nematode Behavior. Cambridge: CABI Publishing; 2004. p. 1–24.

75. Demeure Y, Freckman DW, Van Gundy SD. Anhydrobiotic coiling of nematodes in soil. J Nematol. 1979;11:189–95.

76. Whitford WG, Freckman DW, Elkins NZ, Parker LW, Parmalee R, Philips J, et al. Diurnal migration and response to simulated rainfall in desert soil microarthropods and nematodes. Soil Biol Biochem. 1981;13:417–425.

77. Treonis AM, Wall DH, Virginia RA. The use of anhydrobiosis by soil nematodes in the Antarctic Dry Valleys. Funct Ecol. 2000;14:460–467.

78. de Souza TAJ, de Carli GJ, Pereira TC. Survival potential of the anhydrobiotic nematode *Panagrolaimus superbus* submitted to extreme abiotic stresses. ISJ-Invert Surviv J. 2017;14:85–93.

79. Luan L, Jiang Y, Cheng M, Dini-Andreote F, Sui Y, Xu Q, et al. Organism body size structures the soil microbial and nematode community assembly at a continental and global scale. Nat Commun. 2020;11:6406

80. Stamou GP, Argyropoulou MD, Tsiafouli MA, Monokrousos N, Sgardelis SP, Papatheodorou EM. The study of secondary successional patterns in soil using network analysis: The case of conversion from conventional to organic farming. Pedobiologia. 2011;54:253–59.

81. Creamer RE, Hannula SE, Van Leeuwen JP, Stone D, Rutgers M, Schmelz RM, et al. Ecological network analysis reveals the inter-connection between soil biodiversity and ecosystem function as affected by land use across Europe. Appl Soil Ecol. 2016;97:112–24.

82. Tóth Z, Király I, Mihálka V, Hornung E. Changes in composition and connectivity of soil nematode assemblages under different mulching systems in a strawberry field experiment. Eurasian Soil Sc. 2012;54:1705–1720.

83. Shokoohi, E. Impact of agricultural land use on nematode diversity and soil quality in Dalmada, South Africa. Horticulturae. 2023;9:749.

84. Huang J, Chen J, Huang T, Li G, Wang Z, Zhao S. Soil nematode biodiversity mediates the impact of altered precipitation on dryland agroecosystem multifunctionality in the loess tableland area of China. Agr Ecosyst Environ. 2024;376:109221.

85. McQueen JP, Gendron EMS, Solon AJ, Bueno de Mesquita CP, Hufft RA, Shackelford N, et al. Glyphosate-based restoration of a degraded grassland threatens soil health and the diversity of nematode communities. Soil Biol Biochem. 2024;191:109350.

86. Ptatscheck C, Gansfort B, Traunspurger W. The extent of wind-mediated dispersal of small metazoans, focusing nematodes. Sci Rep-UK. 2018;8:6814.

87. Heyns J. *Elaphonema mirabile n. gen., n. sp*. (Rhabditida), a remarkable new nematode from South Africa. P Helm Soc Wash. 1962;29:128–30.

88. Tschinkel WR. Experiments testing the causes of Namibian fairy circles. PLOS One. 2015;10:e0140099.

89. Jacobson KM, Jacobson PJ. Rainfall regulates decomposition of buried cellulose in the Namib Desert. J Arid Environ. 1998;38:571–83.

90. Jacobson K, van Diepeningen A, Evans S, Fritts R, Gemmel P, Marsho C, et al. Non-rainfall moisture activates fungal decomposition of surface litter in the Namib Sand Sea. PLOS One. 2015;10:e0126977.

91. Kanzaki N, Giblin-Davis RM, Scheffrahn RH, Taki H, Esquivel A, Davies KA, et al. Reverse taxonomy for elucidating diversity of insect-associated nematodes: A case study with termites. PLOS One. 2012;7:e43865.

